# Chemical cues from beetle larvae trigger proliferation and putative virulence gene expression of a plant pathogen

**DOI:** 10.1101/2023.11.21.568124

**Authors:** Marine C. Cambon, Gareth Thomas, John Caulfield, Michael Crampton, Katy Reed, James M. Doonan, Usman Hussain, Sandra Denman, Jozsef Vuts, James E. McDonald

**Affiliations:** School of Natural Sciences, Bangor University, Bangor, UK; School of Biosciences, Institute of Microbiology and Infection, Birmingham Institute of Forest Research, University of Birmingham, Birmingham, UK; Protecting Crops and the Environment, Rothamsted Research, Harpenden, UK; Forest Research, Alice Holt, UK; Department of Geosciences and Natural Resource Management, University of Copenhagen, Rolighedsvej 23, 1958, Frederiksberg C, Denmark

**Author notes:** Corresponding authors: James E. McDonald, Marine C. Cambon, Jozsef Vuts.

## Abstract

Agricultural crop productivity and global forest biomes are coming under increasing threat from insect pests and microbial pathogens. This impact is worsened by inter- kingdom insect-microbe interactions that can increase transmission and disease severity in affected plants. Whilst bacterial chemical cues have been shown to directly influence insect behaviour, the impact of insect-derived compounds on phytopathogens is poorly understood. Here, we investigated the chemical basis for interactions between beetle larvae and bacteria in acute oak decline (AOD), a disease characterised by inner bark necrosis and involving a polymicrobial consortium including *Brenneria goodwinii* and larval galleries of *Agrilus biguttatus*. We found that *A. biguttatus* larval extracts contain chemical elicitors that increase bacterial growth rate and final cell density during *in vitro* culture, and stimulate the differential expression of ∼600 genes, including the Type III Secretion System and its effectors, which are major virulence factors in plant pathogens. Chemical compounds from closely related insect species did not have this effect. These findings highlight the importance of inter-kingdom interactions in plant disease and suggests a novel mode-of-action for insect-derived chemical elicitors in facilitating the virulence of phytopathogens.

## Introduction

Insect-microbe interactions are increasingly important in the context of plant disease owing to increasing incidence of pest and disease outbreaks, driven in part by international trade and climate change (Coolen et al., 2022; Singh et al., 2023). For example, the six-spotted leafhopper *Macrosteles quadrilineatus* is a vector for bacterial phytoplasmas that cause Aster Yellows disease and in-turn benefits from secreted bacterial effectors that targets the plant defence mechanisms (Sugio et al., 2011). Similarly, *Xylella fastidiosa,* which infects a wide range of plant hosts, is transmitted by xylem sap-feeding insects (Sicard et al., 2018). Volatile organic compounds produced by bacteria have been shown to influence insect behaviour (Davis et al., 2013; Weisskopf et al., 2021), which in turn assists in bacterial dispersion (Robacker et al., 2004). A handful of studies have examined the influence of insect metabolites on bacteria, finding inhibitory mechanisms based on the production of antimicrobial compounds (Hall et al., 2011; Teh et al., 2013). However, weather insect-derived compounds have positive positive effects on bacterial activity, increasing growth and/or virulence has not been thoroughly explored. In one example, honeydew produced by aphids has been shown to increase *Pseudomonas syringae* growth (Smee & Hendry, 2022). In addition to serving as a source of nutrition, bacteria use chemical signals as a primary source of information to coordinate their behaviour at the population level (Miller & Bassler, 2001). As such, our limited understanding of the impact of insect-derived compounds on plant-associated bacteria remain a major knowledge gap.Understanding multi-kingdom interactions in plant disease is therefore crucial, especially in decline diseases such as Acute Oak Decline (AOD), where insects could play a pivotal role. AOD is a complex decline disease affecting native British oak species (*Quercus robur* and *Quercus petraea*) and recently identified in many European countries (Carluccio et al., 2024; Ruffner et al., 2020; Tkaczyk & Sikora, 2024). It involves the cumulative action of multiple biotic and abiotic factors, such as larvae of bark-boring beetles and bacterial phytopathogens (Denman & Webber, 2009). The disease is characterised by inner bark tissue necrosis caused by a polymicrobial consortium composed of at least three bacterial species (*Brenneria goodwinii*, *Gibbsiella quercinecans*, and *Rahnella victoriana*), and characterised by the presence of larval galleries of the two-spotted oak buprestid, *Agrilus biguttatus* on the phloem–sapwood interface.

*B. goodwinii* is recognised as the dominant pathogen within necrotic lesions (Denman et al., 2018) and transcriptomic analysis of *B. goodwinii* in oak stem tissue revealed the up- regulation of putative virulence genes (Doonan et al., 2020). Co-infection of *A. biggutatus* larvae significantly increased the expression of these genes including Type III secretion system effectors *srfB* and *avrE,* known to contribute to virulence and suppression of host immunity in other pathosystems (Worley et al., 2000) (Degrave et al., 2015), and *katG* which detoxifies hydrogen peroxide released as part of the host defence response (Jittawuttipoka et al., 2009). The presence of *A. biguttatus* larvae also increased the area of necrotic tissue caused by bacteria (Denman et al., 2018; Doonan et al., 2020). Taken together, these findings suggest that the interaction between *B. goodwinii* and *A. biguttatus* larvae in the inner bark enhances bacteria-induced necrosis and facilitates the proliferation and spread of damaging bacteria within infected trees.

Although interactions between *B. goodwinii* and *A. biguttatus* larvae have been demonstrated *in planta* (Denman et al., 2018; Doonan et al., 2020), the mechanism underpinning this interaction remains unknown. In this study, we inoculated oak logs with *A. biguttatus* eggs and collected late-instar larvae and gallery wood tissue from the logs. As *A. biguttatus* larval feeding may chemically modify their environment and/or trigger host responses, we extracted chemicals from asymptomatic as well as gallery wood tissue and the bodies of *A. biguttatus* larvae and independently assessed their bioactivity towards *B. goodwinii* in controlled *in vitro* experiments. We tested three hypotheses: (1) *B. goodwinii* proliferation and virulence gene expression are indirect effects of *A. biguttatus* larval activity and are triggered by chemicals associated with gallery wood tissue; (2) *B. goodwinii* proliferation and virulence gene expression are direct effects of *A. biguttatus* larvae and are triggered by larval chemical elicitors; (3) the interaction between *B. goodwinii* and *A. biguttatus* larvae is specific to *A. biguttatus*. We show that chemical elicitors from *A. biguttatus* larvae (hypothesis 2), but not gallery wood tissue (hypothesis 1), directly trigger an increase in *B. goodwinii* growth and the up- regulation of 307 genes, including effectors involved in bacterial Type III secretion system. Such an effect was not observed with extracts from closely related wood-boring beetle species, suggesting some specificity in this interaction (hypothesis 3). Determining the mechanisms of insect-microbiota interactions and their emergent impacts on plant disease outcomes is critical to inform integrated pest management strategies for globally important plant diseases.

## Results

### Gallery wood tissue extracts do not increase bacterial proliferation compared to healthy wood tissue extracts

To test the effect of chemical extracts from galleries made by *A. biguttatus* larvae on *B. goodwinii* proliferation (hypothesis 1), we compared bacterial growth in cultures supplemented with asymptomatic wood extract and gallery wood tissue extract (Figure 1A). At 24 hpi (hours post inoculation), cultures reached an OD600 of 0.75 ± 0.05, 0.78 ± 0.09 and 0.9 ± 0.06 for asymptomatic wood tissue extracts, gallery wood tissue extracts and water control, respectively (Figure 2A). The growth rate of cultures supplemented with gallery wood extracts (0.30 ± 0.04) was significantly lower than those supplemented with asymptomatic wood extracts (0.39 ± 0.03, Figure 2B, pairwise Wilcoxon test, p-value < 0.05). The CFU density at 24 hpi did not differ significantly between gallery wood tissue and asymptomatic wood treatments (12.69 ± 2.5 x 10^8^ CFU/mL and 12.32 ± 10.0 x 10^8^ CFU/mL, respectively Figure 2C) suggesting that *B. goodwinii* grows faster with asymptomatic wood extract but the number of living cells in stationary phase is the same for both asymptomatic and gallery wood extracts. These results show that chemicals compounds extracted from gallery wood tissue do not cause more proliferation of *B. goodwinii* than asymptomatic wood chemicals compounds.

**Figure 1:**
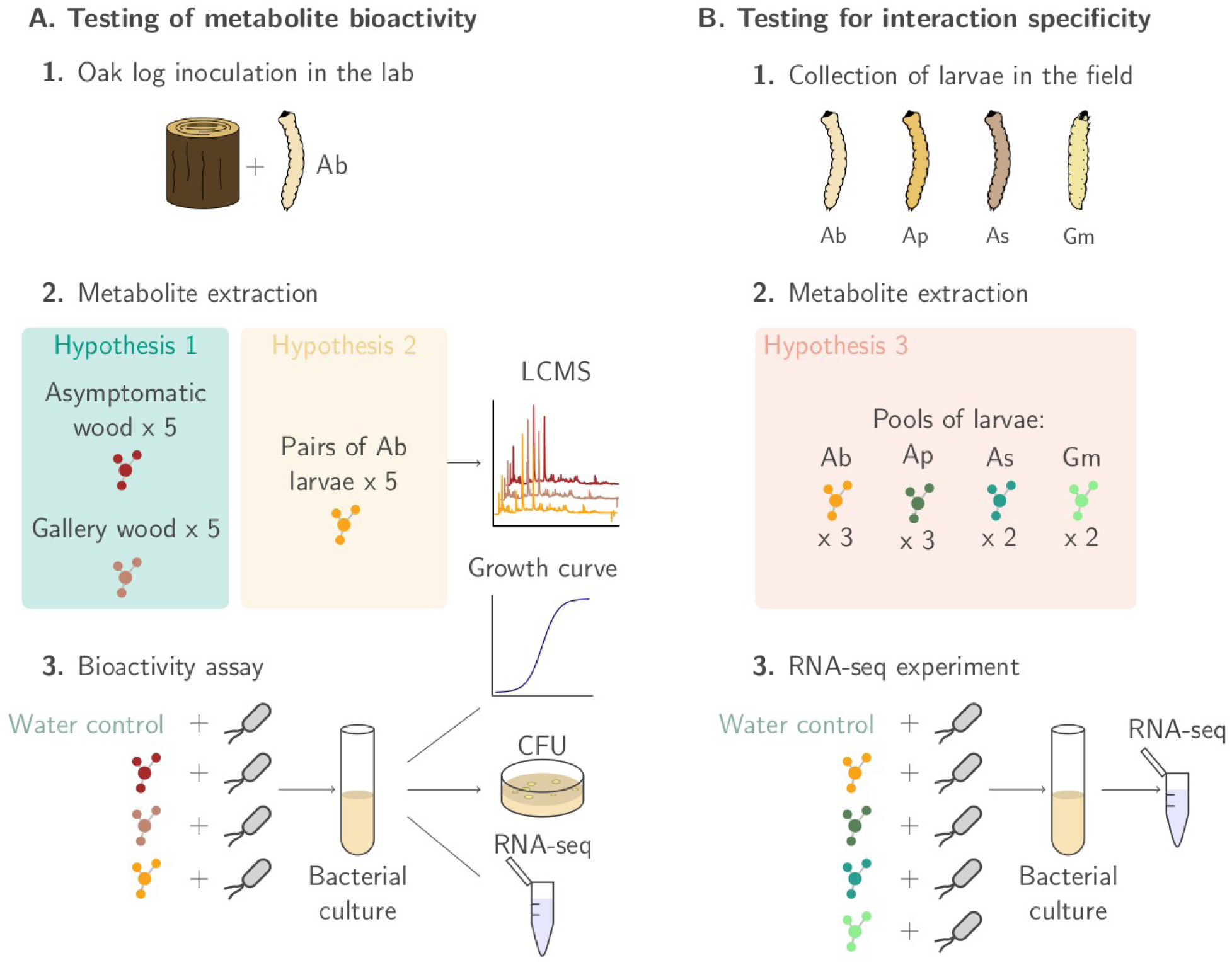
Summary of the experimental design. **A.** For hypotheses 1 and 2, oak logs were inoculated in the lab to yield Ab larvae and wood tissue samples for extraction. **B.** For hypothesis 3, *Agrilus* larvae (Ab, Ap and As) were collected from the field or purchased (Gm). Metabolites from the larvae were extracted and tested for bioactivity on *Brenneria goodwinii* cultures. Ab = *Agrilus biguttatus*; Ap = *A. planipennis*; As = *A. sulcicollis*; Gm = *Galleria mellonella*

**Figure 2:**
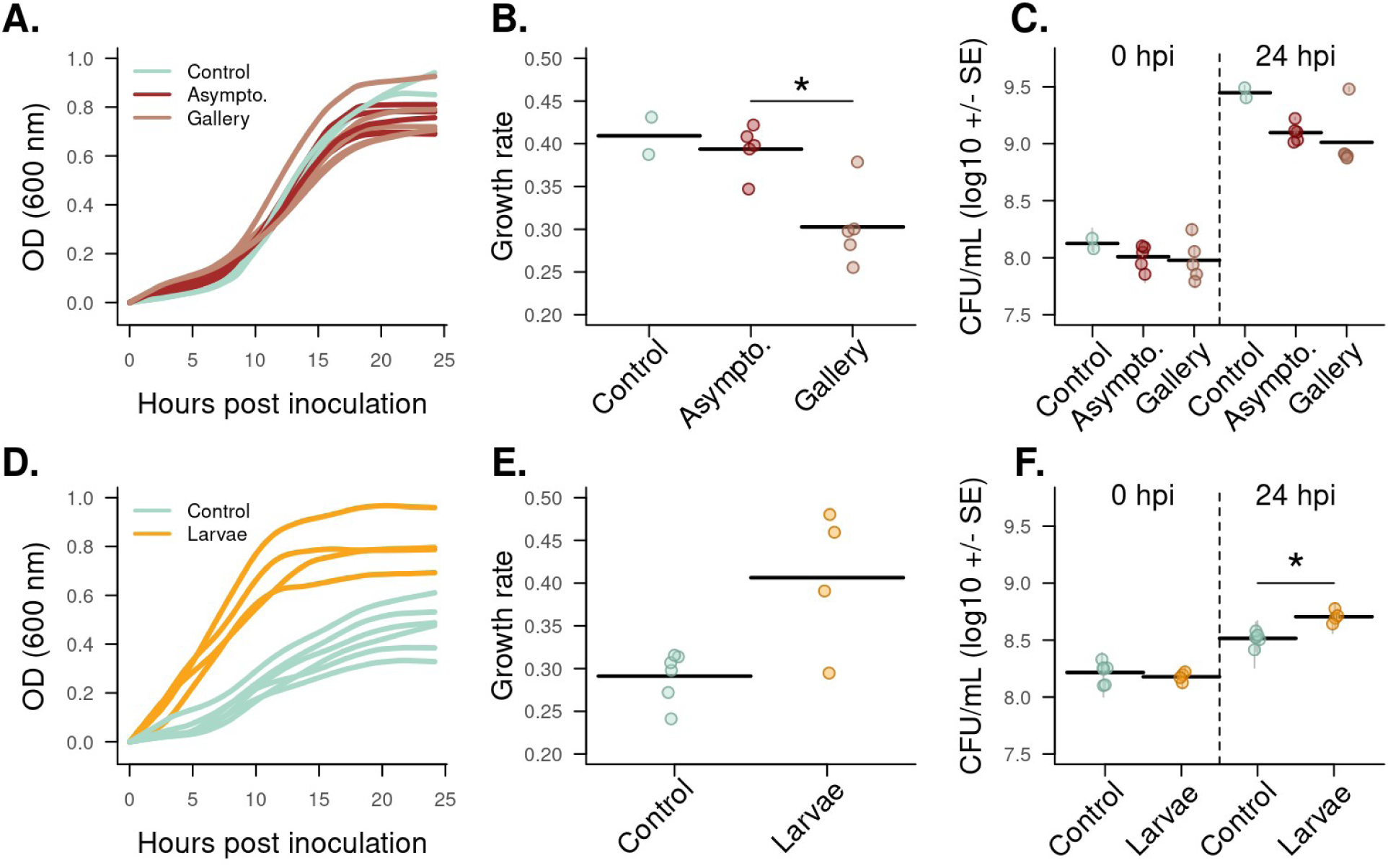
***B. goodwinii* growth curves (A), growth rates (B), and cell density (C)** obtained in 3 mL liquid cultures supplemented with gallery wood tissue (Gallery) and asymptomatic wood (Asympto.) chemical compounds, compared to chemical compound extraction blanks (Controls). Growth curves (D), growth rates (E), and cell density (F) obtained in 200 µL liquid cultures supplemented with beetle larvae body extracts (Larvae), compared to chemical compound extraction blanks (Controls). The controls were the same in both experiments but the difference in growth rate observed in (A) and (D) are due to the different volumes of cultures. An additional experiment replicating the results of (A) in the same volume as (D) can be found in Figure S1. Stars indicate significant differences (pairwise Wilcoxon test (above) or t- test (below), p-value < 0.05). OD = optical density; CFU = colony-forming unit; SE = standard error

### *A. biguttatus* larval elicitors cause bacterial proliferation

To test the direct effect of *A. biguttatus* (Ab) larval chemical elicitors on *B. goodwinii* proliferation (hypothesis 2), we compared bacterial growth in cultures supplemented with Ab larval extract or water (extraction blank control). At 24 hpi, cultures in a 200 µL volume reached an OD600 of 0.75 ± 0.09 and 0.47 ± 0.1 for Ab extracts and water control, respectively (Figure 2D). The growth rate of cultures supplemented with Ab extracts was higher than that of control cultures (0.41 ± 0.07 and 0.30 ± 0.04, respectively; Figure 2E). This trend was confirmed by the Ab larval bulk extract growth curves in Figure 4B. The CFU density at 24 hpi was also significantly higher for cultures supplemented with Ab extract than for control cultures (5.1 ± 0.67 x 10^8^ CFU/mL and 3.3 ± 0.4 x 10^8^ CFU/mL, respectively; Figure 2F). These results indicate that chemical elicitors extracted from *A. biguttatus* larvae stimulate bacterial growth.

### *A. biguttatus* larval elicitors modify bacterial gene expression profile and induce putative virulence gene up-regulation

To test for the impact of chemical elicitors on bacterial gene expression, we compared the gene expression profile of *B. goodwinii* in cultures supplemented with asymptomatic and gallery wood tissue extracts (hypothesis 1), and *A. biguttatus* larval extracts and water control (hypothesis 2). Cultures supplemented with gallery wood tissue or asymptomatic wood extracts did not have a clearly distinct gene expression profile and did not cluster separately based on the correlation matrix of normalised transcript count per million (CPM) (Figure 3A). A total of 12 genes were differentially expressed between the gallery and asymptomatic wood tissue, with 3 up-regulated and 9 down-regulated genes in the gallery wood tissue treatment (Table S1). Two out of the 3 up-regulated genes were linked to flagellum production (*fliA*, *fliF* Figure 4). These results show that chemical compounds extracted from gallery wood tissue have a small impact on the gene expression profile of *B. goodwinii* compared to extracts from asymptomatic wood tissue and mainly impact flagellar genes.

**Figure 3:**
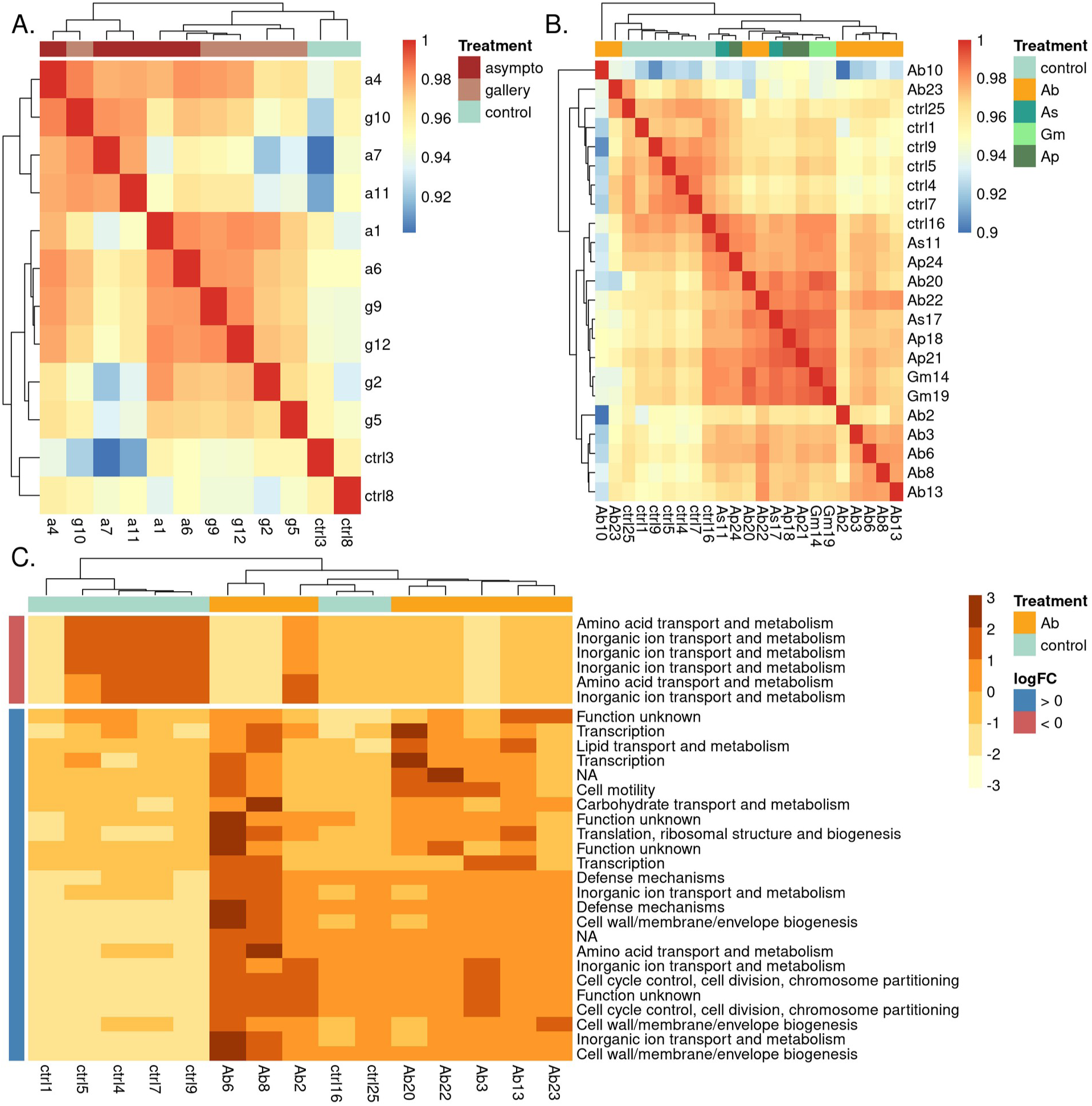
Gene expression profiles obtained from RNAseq data of *B. goodwinii* cultures. supplemented with chemical extracts from oak wood tissues, insect larvae or water (control). **A.** and **B.** Heatmaps of the sample correlation matrix based on the normalised transcript counts per million (CPM). Asympto = asymptomatic wood extract, gallery = gallery wood tissue extract, Ab = *A. biguttatus* larval extract, Ap = *A. planipennis* larval extract, Gm = *Galleria mellonella* larval extract. Sample Ab10 had a gene expression profile different from all other samples with low correlation values and was considered as an outlier and removed from the differential expression analysis. **C.** Heatmap of the centred and scaled normalised transcript CPM of the top 30 differentially expressed genes in Ab treatment, compared to control. Row names correspond to the Clusters of Orthologous Groups of proteins (COGs) annotation for each gene.

**Figure 4:**
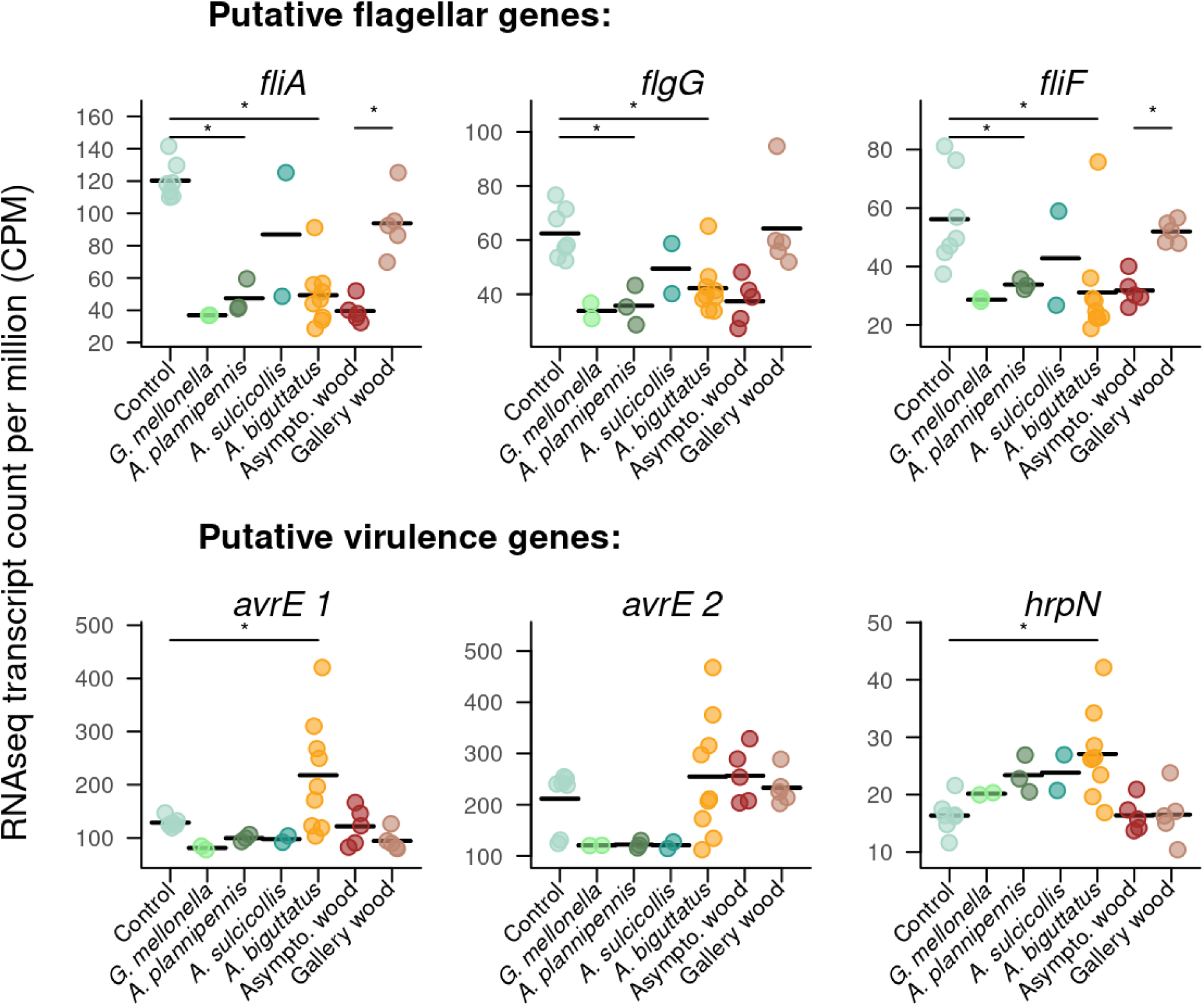
RNAseq transcipt CPM for genes of interest. These include putative flagellar and virulence genes. Bars with stars represent significantly differentially expressed genes based on the edgeR differential expression analysis.

In contrast, cultures supplemented with *A. biguttatus* larval extracts had a distinct gene expression profile and formed a separate cluster from the control cultures (no larval extract) based on the correlation matrix of normalised transcript CPM (yellow and light blue samples in Figure 3B). A total of 585 genes were differentially expressed between *B. goodwinii* cultures receiving Ab larval extracts and the control treatment, with 307 up- regulated and 278 down-regulated genes in the Ab larvae treatment (Table S2). The 30 most differentially expressed genes and their functional annotation are represented in Figure 3C. Up-regulated genes included the Type III secretion system apparatus and effectors (*avrE*, *hrpN,* Figure 4) which have been found to be expressed during lesion formation *in planta* (Broberg et al., 2018; Doonan et al., 2020). These results show that Ab larval elicitors have a significant impact on bacterial gene expression and trigger the expression of putative bacterial virulence genes.

### Extracts from other beetle species influence *B. goodwinii* gene expression but do not induce putative virulence genes up-regulation

To test for the specificity of the bacteria-beetle interaction in AOD, we replicated the RNAseq experiment using two other species of *Agrilus*: the European oak borer *A. sulcicollis* (As) and the emerald ash borer *A. plannipenis* (Ap). We also included in the experiment the wax moth *Galleria mellonella* (Gm) as an outgroup. Because of the limited availability of larvae extract, only one experiment was performed and culture were grown for 10 hpi only to allow sampling for RNAseq analysis. OD600 measurements from these 10 first hours of growth were performed on some of the cultures and show a trend of higher growth rates for Ab, Ap, and Gm compared to As and control (Figure S2). However, differences in growth rates were not significant which may be due to the small sample size for these measurements.

*B. goodwinii* cultures supplemented with larval extracts clustered together and away from the water control based on the correlation matrix of normalised transcript CPM (Figure 3B). However, the different species of larvae did not necessarily form distinct species-specific clusters, indicating some variability in the bacterial response to larval chemical compounds. Nevertheless, most Ab samples clustered separately from the other larval extracts. Cultures supplemented with As and Gm extracts did not show any significant differentially expressed genes compared to the control, although data shows a trend of higher CPM for *hrpN* compared to control. Cultures supplemented with Ap extracts showed 125 differentially expressed genes, including 85 up-regulated and 40 down-regulated genes (Table S3). Most up-regulated genes were related to cell multiplication, while down-regulated genes contained 9 flagellar production- and motility-related genes. However, the putative virulence genes that were differentially expressed in the presence of Ab larval extracts were not significantly differentially expressed in the presence of Ap larval extracts, although *hrpN* shows a trend of higher CPM compared to the control (Figure 4).

### Bacterial proliferation is triggered by a fraction of the *A. biguttatus* larval elicitor extract

To narrow down identification of the bioactive fraction(s) of the *A. biguttatus* larval extracts, we compared *B. goodwinii* growth in cultures supplemented with the bulk larval extract with that of cultures supplemented with four fractions of the bulk larval extract (Fraction 1 (F1) to Fraction 4 (F4); Figure 1B and 5A). At 24 hpi, cultures supplemented with the bulk larval extract, F1 or F4 reached greater OD600 (0.50±0.04, 0.51±0.05 and 0.48±0.01, respectively; Figure 5B) compared to cultures supplemented with water (control), F2 or F3 (0.36±0.03, 0.38±0.03 and 0.40±0.05, respectively; Figure 5B). The growth rate of *B. goodwinii* in cultures supplemented with F1 was significantly higher than any other treatments, including the bulk larval extract. The growth rate of cultures supplemented with F2, F3 and F4 did not significantly differ from the water control (Figure 5C). The cell density at 24 hpi was significantly reduced for *B. goodwinii* in cultures supplemented with F1 and F4 compared to the other treatments (Figure 5D). Altogether, these results suggest that F1 contains the chemical elicitors responsible for the proliferation (higher growth rate) of *B. goodwinii*, but that these chemical compounds, when separated from the other fractions, reduce cell density in the stationary phase of growth.

**Figure 5:**
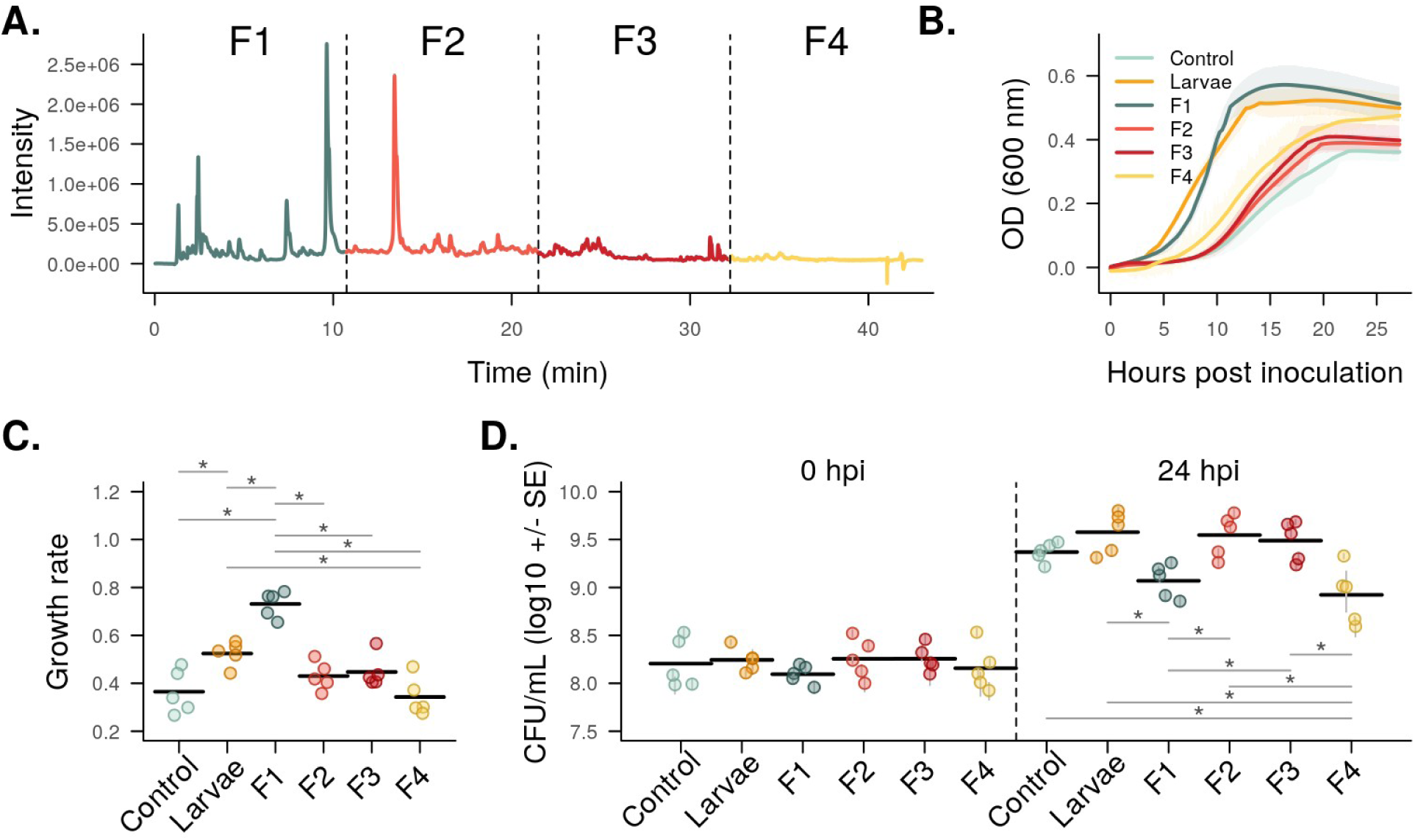
Bioactivity of different fractions of the *A. biguttatus* larval extract. **A.** LC-MS chromatogram of the bulk larval extract (“Larvae” treatment on panel B to D). A part of it was then fractionated in four fractions (fraction F1 to F4 on panel B to D). **B.** Growth curves, **C.** growth rates, and **D.** cell density were obtained in 200 µL cultures supplemented with the bulk larval extract (Larvae), the four larval extract fractions (F1 to F4) and water (Control). Bars with stars indicate significant differences (Tukey HSD, p-value < 0.05). OD = optical density; CFU = colony-forming unit; SE = standard error

## Discussion

Investigations of insect-bacteria interactions in plant diseases have largely focussed on the role of bacterial volatile organic compounds (VOCs) in modulating insect behaviour (Davis et al., 2013; Leroy et al., 2011; Weisskopf et al., 2021). Less attention has been paid to the influence of insect-derived chemical cues on bacterial growth and virulence. In this study, we investigated the impact of insect chemical elicitors on a bacterial species associated with Acute Oak Decline (AOD). We found that *A. biguttatus* larval elicitors, but not gallery wood tissue extracts, increase *B. goodwinii* proliferation and virulence gene expression. Elicitors from some other insect species influenced bacterial gene expression but did not trigger the expression of T3SS effector genes.

Extracts from larval gallery wood tissue did not cause bacterial proliferation as compared to healthy wood controls, suggesting that proliferation of the bacteria around *A. biguttatus* galleries *in planta* (Denman et al., 2018) is not mediated by the tree host chemicals. The growth rate of cultures was in fact lower for extracts from gallery wood tissue compared to healthy wood tissue; potentially owing to tree host defence compounds produced in response to larval invasion (Broberg et al., 2018). However, the final cell density was not significantly different between gallery and healthy wood tissue extract treatments, suggesting that the extracted chemicals only affect the speed of proliferation but not final cell density. In addition to a reduction of growth rate, the wood gallery extracts triggered the expression of flagellum-associated genes, which are mainly involved in bacterial motility (Thomas et al., 2001; Wilkinson et al., 2011) and could represent a mechanism for the bacteria to move away from an unsuitable environment. Some flagellar genes are involved in virulence and plant wall adherence in certain plant pathogens (Jahn et al., 2008; Malamud et al., 2011) which could indicate the activation of some virulence process components. However, any such indirect effect of *A. biguttatus* larvae on *B. goodwinii* virulence gene expression through the modification of the wood tissue environment is thus less powerful than the direct effect of larval extracts but cannot be completely excluded.

*A. biguttatus* larval extracts increased bacterial proliferation *in vitro,* suggesting that the interaction between insect and bacteria observed *in planta* is at least in part directly mediated by larval chemical compounds. Previous studies conducted on artificially infected oak logs showed that the lesion size caused by AOD bacteria significantly increased in the presence of beetle larvae (Denman et al., 2018). In addition, AOD lesions are observed in close proximity to beetle galleries *in natura,* which suggests a bacteria-beetle interaction, and/or the dissemination of the bacteria within wood tissues by the beetle larvae. In the light of our results, we hypothesise that elicitors are released by *A. biguttatus* larvae as they form galleries, and that these chemical compounds trigger the proliferation of *B. goodwinii*, which starts degrading the wood. However, we cannot exclude the possibility that elicitors may be metabolised by the bacteria and may act as a food source in our experiment.

The link between larval elicitor production and bacterial lesion formation is further supported by the fact that *A. biguttatus* larval extracts also triggered the expression of putative virulence genes in *B. goodwinii.* Among the 307 significantly up-regulated genes induced by larval extracts were some potential virulence genes associated with a Type III secretion system (T3SS), which is well reported in other plant pathogens. These include the *hrpN* gene encoding for the harpin component of the needle structure of the T3SS in *Pseudomonas syringae* (Collmer et al., 2000) and has been found over-expressed in AOD lesions *in situ* (Broberg et al., 2018). This T3SS belongs to the Hrc-Hrp1 family only present in plant pathogens and is distinctive in that it possesses a long, flexible pilus able to traverse the thick plant cell wall, suggesting that the target of this T3SS is the plant host (Troisfontaines & Cornelis, 2005). In addition, we found a*vrE* genes up-regulated in the presence of *A. biguttatus* larval elicitors, which were previously reported to be up- regulated in the presence of *A. biguttatus* larvae *in planta* (Doonan et al., 2020). The a*vrE* gene encodes for a T3SS effector involved in the suppression of plant immunity by *P. syringae* and other plant pathogens (Degrave et al., 2015) which induce water flux into host cells resulting in cell swelling and bursting (Nomura et al., 2023). These results give another strong indication that beetle larvae are involved in triggering the virulence of *B. goodwinii.* It should be noted, however, that some bacteria have also been reported to possess different T3SS effectors to allow interactions both with plant hosts and insect vectors (Correa et al., 2012). The response of *B. goodwinii* to *A. biguttatus* chemical compounds could thus potentially be involved in the colonisation of larvae by the bacteria; a*vrE* gene knock-outs could confirm its function in the AOD system.

We investigated the specificity of this insect-bacterium interaction by extracting chemical compounds from the European oak borer *A. sulcicollis*, the emerald ash borer *A. planipennis*, and the wax moth *Galleria melonella.* We did not find any significant differentially expressed genes in *B. goodwinii* cultures supplemented with *A. sulcicollis* and *G. mellonella* compared to the control, which could be explained by the lower sample size we had for these insect species. However, *B. goodwinii* cultures supplemented with *A. planipennis* extracts showed 125 differentially expressed genes compared to the water control. Among these, numerous motility-related genes were down-regulated, which is similar to the response of *B. goodwinii* to *A. biguttatus* extracts and indicate a change of phenotype in response to insect-derived chemical compounds. However, none of the T3SS-related genes found up-regulated in the presence of *A. biguttatus* were differentially expressed in response to *A. planipennis* extracts. These results suggest that chemical compounds from different insect species can show bioactivity towards *B. goodwinii,* but the trigger of virulence gene expression is specific to *A. biguttatus* under the conditions of our experiment.

One fraction of the *A. biguttatus* larval extract produced the same growth curves as the bulk larval extract when added to a culture of *B. goodwinii*. The growth rate obtained with this fraction was in fact higher than with the bulk larval extract, while the final cell density at 24 hpi was lower. This indicates that several chemical compounds are necessary to reproduce the full phenotype observed when supplementing cultures with the bulk larval extract, which may be present in several of the four fractions. Identifying the bioactive chemical compounds should be a future priority but it is hindered by the limited availability and seasonality of *A. biguttatus* larvae and the predicted significant challenges associated with the identification work. However, we successfully narrowed down the activity range of separate fractions of larval extracts responsible for the proliferation and up-regulation of effector gene expression in *B. goodwinii,* which will aid chemical compound characterisation efforts.

The strong effects of *A. biguttatus* extracts on *B. goodwinii* in comparison to those from gallery wood tissue strengthens the argument that *A. biguttatus* acts as a synergistic factor with *B. goodwinii*, the major pathogen found in AOD lesions. Whether the bacteria are present in the tree in a non-pathogenic state before beetle invasion, or if they are vectored by the beetle to the tree remains to be answered, but our study emphasises the necessity of integrating *A. biguttatus* population survey and management into programs tackling AOD. The potential benefits this interaction provides to each partner need to be further evaluated to understand their evolutionary consequences. It is possible that this interaction results from co-adaptation and provides a benefit to *A. biguttatus* beetle larvae, as *B. goodwinii* expresses detoxification genes when infecting the tree (Denman et al., 2018; Doonan et al., 2020). This may provide protection for the larvae against plant defences as well as weakening the tree, thus increasing larval survival and overall reproductive success. For the bacteria, supporting the growth and emergence of beetle larvae could potentially lead to facilitated multiplication and transmission between trees. On the other hand, this interaction could equally be incidental and not a result of any evolutionary relationship, with no impact on the bacteria or beetle fitness.

Although *A. sulcicollis* shares its oak hosts with *A. biguttatus,* the same interaction with *B. goodwinii* was not observed in our experiment. Anecdotally, *A. sulcicollis* colonises very poorly defended or recently dead oak tissues, which are drier and more degraded than those colonised by *A. biguttatus* (K. Reed, personal observation). Perhaps it therefore does not have the same need for protection from host defences as *A. biguttatus,* which is able to colonise stressed, but still living host tissue. On the other hand, *A. planipennis,* a highly aggressive species, which is able to kill healthy ash trees, did provoke differential gene expression in *B. goodwinii.* Given the economic cost of damage due to *Agrilus* pests, future studies should explore the interaction between Agrilus species and microbial communities in trees.

Although the importance of interactions between insects and bacteria is well established in the case of symbiosis (Douglas, 2009) or insect behaviour manipulation by bacterial VOCs (Davis et al., 2013), our suggests another form of insect-bacteria interaction, where insect-derived chemicals modulate bacterial gene expression potentially influencing virulence. The implications of this finding are of particular importance in the context of recent increases in plant disease outbreaks involving pest-pathogen interactions. Our study together with previous work strongly suggest that an insect herbivore affects bacterial behaviour by triggering the expression of virulent phenotypes and facilitating lesion formation in live bark tissue, thus contributing to tree decline (Broberg et al., 2018; Doonan et al., 2020). Such overlooked interactions warrant future investigation to safeguard forest biomes.

## Material and methods

### Gallery wood tissue and larvae sample production

Sections of bark and sapwood cut from *A. biguttatus* (Ab) infested oak trees were stored in purpose-built, walk-in emergence cages at Forest Research (Farnham, UK). After the adults emerged, they were collected and maintained in the lab as described by (Reed et al., 2018). A maximum of 8 beetles were housed per 2 L cage, and the beetles were provided with freshly-cut oak leaves and 20 % sugar solution and water on cotton wool. Beetle eggs, laid on paper towels placed underneath the cages, were collected twice weekly and incubated in 15 mL Falcon tubes at 17.5 °C until 1 or 2 days before the hatch time predicted by the egg hatching model (Reed et al., 2018). Oak billets were cut to a length of approximately 90-100 cm and allowed to stand in water for 2 weeks prior to inoculation. 10-15 eggs were placed in wounds made in the phloem with a 5 mm cork borer (4 wounds per billet). Infested billets were incubated at room temperature for 3 months before harvesting of Ab larvae from the billets. Ab larvae and wood tissue samples were collected by removing the outer bark with a wood knife to expose larval galleries. Undamaged Ab larvae were collected in 1.5 mL tubes and frozen into liquid Nitrogen. Asymptomatic wood tissue and gallery wood tissue were collected in 15 mL tubes and frozen in liquid Nitrogen. Samples were stored at -80 °C until chemical extraction.

Final instar larvae of and *Agrilus sulcicollis* (As) were collected from the phloem of recently-dead, infested large oak branches (∼15cm diameter) and stems at Richmond Park, Greater London. They were morphologically differentiated from *A. biguttatus,* which is the only other species found in large oak branches and stems at Richmond Park, using the key in (Volkovitsh et al., 2020).

Third and fourth instar *Agrilus planipennis* (Ap) larvae were produced on *Fraxinus excelsior* billets within the Holt Containment Facility at Forest Research, Farnham, as follows. Eggs, laid onto filter paper, were couriered from the USDA APHIS PPQ EAB Biocontrol Facility (Brighton, UK). 8 eggs per billet were cut out from the filter paper and wrapped with Parafilm to hold them snugly against the bark. The billets were kept standing in trays of water at 25 °C for 3-4 weeks before the bark was peeled and the larvae were harvested. After collection, undamaged Ab, As and Ap larvae were frozen in 15 mL tubes at -80 °C, where they were stored until processing. Finally, *Galleria melonella* (Gm) larvae were commercially purchased (Livefood UK Ltd) and stored at -80 °C until processing.

### Larval and wood tissue chemical compound extraction and analysis

#### Chemical extraction

Wood tissue chemical compounds were extracted from 2 x 2 cm (64±20 mg) sections of inner bark material (either asymptomatic or containing larval galleries) in 5 mL 100 % MeOH for 2 h. To remove cellular debris, extracts were filtered through a plug of silanized glass wool held within a Pasteur pipette, into pre-weighed glass vials. Resultant extracts were dried to completion under a gentle stream of N2. Five biological replicates were performed per treatment, and an extraction blank containing only 100 % methanol was also included.

Ten snap-frozen Ab larvae collected from the oak log trial were weighed and individually extracted in 5 mL of 100 % methanol for 2 h. To remove cellular debris, extracts were filtered through a plug of silanized glass wool held within a Pasteur pipette, into pre- weighed glass vials. Five extraction blanks containing only 100 % methanol were also included. 400 µL of each extract were kept for LC-MS (liquid chromatography and mass spectrometry) analysis. The remaining of the extracts were dried to completion under a gentle stream of N2. Dried extracts were resuspended in Milli-Q water to a 3 mg/mL concentration and filter sterilised through a 0.22 µm pore size filter, for bioactivity tests. The 10 larval extracts were then pooled in pairs in equal volume to obtain sufficient amount of chemical compounds for bioactivity tests, representing five biological replicates, and stored at -20 °C until processing.

Chemical compounds were extracted from As larvae, Ap larvae and Gm larvae following the same protocol. Due to different quantities of chemical compounds extracted from different species, larval extracts were pooled, e.g. two or three individual larvae were pooled to obtain at least 3 biological replicates per insect species at 3 mg/mL.

#### LC-MS conditions

LC-MS analysis was carried out using an Acquity ultra-high-pressure liquid chromatography (UPLC) system coupled to a Synapt G2Si Q-Tof mass spectrometer with an electrospray ionisation source (Waters, UK). The system was controlled through Masslynx 4.1 software (Waters). Chromatographic separation was carried out at a flow rate of 0.21 mL.min^-1^ using a UPLC BEH C18 column (2.1 x 150 mm, 1.7 μm, Waters) coupled to a C18 Vanguard pre-column (2.1 x 5 mm, 1.7 μm, Waters). The mobile phase consisted of solvent A (0.01% formic acid v/v, water) and solvent B (0.01% formic acid v/v in methanol) with the following gradient: 0 min, 95:5 (A:B), to 85:15 at 7 min, 75:25 at 11 min, 70:30 at 13 min, 50:50 at 18 min; 25:75 at 25 min; 0:100 at 30 min, and 95:5 at 39.1 min.

An Acquity photodiode array (PDA) detector was used to monitor the UV trace (range 200-450nm), sampling rate of 10 points s-1 with resolution set to 2.4 nm. The Synapt was operated in sensitivity mode and set to a mass range of 50 – 1200 Da and scan time = 0.1 s, in both ionisation modes. The system was operated in MS1 mode with the following conditions: capillary voltage – 2.5 KV, sample cone voltage 30 V, sample offset 80 V, source temperature 100 °C, desolvation temperature 300 °C, desolvation gas flow 800 L.h^-1^, cone gas flow 57 L.h^-1^.

#### Preparative HPLC conditions

To locate active fractions within the larval extracts, preparative HPLC was performed by repeated injections on a Hichrom ACE 5 C18 column (250 × 4.6 mm i.d.) connected to Shimadzu Nexera UC SFC-HPLC hybrid system. The mobile phase consisted of solvent A (water) and solvent B (methanol) with the following gradient: 0 min, 95:5 (A:B), to 85:15 at 7 min, 75:25 at 11 min, 70:30 at 13 min, 50:50 at 18 min; 25:75 at 25 min; 0:100 at 30 min, and 95:5 at 39.1 min. The injection volume was 100 µL (five successive injections), and the separation was carried out over 43 min with a flow rate of 1 mL.min^- 1^. Fractions were collected every 10.75 min, yielding four individual fractions. The four fractions collected using prep-HPLC were evaporated to 1 ml using a rotary evaporator, after which the fractions were dried under a gentle stream of nitrogen.

### Chemical compound bioactivity analysis

#### Bacterial strain and medium

Bioactivity assays were conducted on the type strain *Brenneria goodwinii* FRB141 kept in glycerol stocks at -80 °C. Before the experiment, the strain was streaked onto Nutrient Agar (LAB M Limited) and incubated for 48 h at 28 °C. Starter cultures were then prepared by picking a single colony and growing it overnight in 10 mL of Nutrient Broth n°2 (Oxoid) at 28 °C and 150 rpm shaking. Starter cultures were rinsed twice by pelleting the cells for 5 min at 3000 g, discarding the supernatant and adding 10 mL of ¼ strength Ringers solution (LAB M Limited). Cells were then pelleted again, and OD (Optical Density) at 600 nm was adjusted to 0.5 in ¼ strength Ringers solution.

Subsequent growth measurements were performed in M9+glu minimal medium composed of M9 salt (Na2HPO4 42 mM; KH2PO4 22 mM; NaCl 8.55 mM; NH4Cl 18.6 mM; MgSO4•7H2O 2 mM; CaCl2 0.1 mM) and 20% glucose, with or without the addition of chemical compounds tested for bioactivity.

#### Growth experiment

The bioactivity test of gallery wood tissue chemical compounds was performed in culture broth composed of 1 mL of gallery wood tissue or asymptomatic wood tissue extract (3 mg/mL), 0.66 mL of M9+glu minimum medium (4.5 x concentration), 1.04 mL of water, and 0.3 mL of starter culture (starting OD600 of 0.05), for a final volume of 3 mL. For controls, wood tissue extracts were replaced by chemical compound extraction blanks. Chemical compounds from 5 gallery wood tissue samples and 5 asymptomatic wood samples were tested (5 biological replicates per group and 1 technical replicate) and cultures were placed in a 12-well plate. To allow for comparison with hypothesis 2 which could not be performed in a 3 mL volume due to the lower quantity of larval chemical compounds obtained from the extraction step, we replicated the growth experiment in a 200 µL volume (Figure S1).

Because the number of Ab larvae available for the experiment was limited, the quantity of chemical extract available for the experiments was small. Consequently, the test of Ab larval chemical compounds on bacterial growth was performed in a smaller culture volume. The broth was composed of 60 µL of larval extract (3 mg/mL), 40 µL of M9+glu minimum medium (4.5X), 80 µL of water, and 20 µL of starter culture (starting OD600 of 0.05), for a final volume of 200 µL. For controls, Ab larval extracts were replaced by chemical compound extraction blanks. Chemical compounds from 5 pairs of Ab larvae were tested with 6 technical replicates to account for the higher noise in OD measurement when performed in small culture volumes and cultures were placed in a 96-well plate.

For each growth experiment, a replicate plate containing the same liquid growth medium, but without bacterial inoculation, was performed to get OD background correction with sterile medium and check for medium contamination. Plates were placed into the Stratus plate reader (Cerillo) for OD600 measurement every 3 minutes over the course of the experiment. The plate reader was incubated at 28 °C and shaken at 150 rpm for 24 h. A 20 µL aliquot of each culture was collected at 0 hpi (hours post inoculation) and 24 hpi for Colony Forming Unit (CFU) quantification.

The number of viable cells in the cultures was estimated at 0 hpi and 24 hpi by adding 20 µL of culture to 180 µL of sterile ¼ strength Ringers solution and completing a serial dilution in the same way up to 10^-8^ in a 96-well microplate. Six replicates of 10 µL of each dilution were then pipetted onto Nutrient Agar and incubated for 24 h at 28 °C.

The number of CFUs was counted from the appropriate dilution, and the total CFU/mL of culture was estimated as [inline] concentration from the 6 replicate 10 µL drops were averaged and standard error was calculated.

#### Transcriptome experiments

To ensure sufficient biomass for RNA-seq analysis, cultures were set up in a final volume of 3 mL as described above, and larval extracts had to be pooled in pairs to obtain enough material for RNA extraction. Three separate RNA-seq experiments were performed with the following sample sizes: Experiment 1: extracts from 5 gallery wood tissue samples and 5 asymptomatic wood tissue samples; Experiment 2: extracts from 5 pairs of Ab larvae and 5 extraction blanks; Experiment 3: extracts from 3 pairs of Ab larvae, 3 pools of Ap larvae, 2 pools of As larvae, 2 pools of Gm larvae and 3 extraction blanks. Extracts were added to culture broth as described above and culture plates were incubated for 10 hours to obtain cultures in mid-exponential phase of growth. Cultures were transferred into 2 mL screw cap tubes, centrifuged for 5 min at 3000 rcf to discard the supernatant, and immediately frozen into liquid nitrogen. Cell pellets were then stored at -80 °C until RNA extraction.

The total RNA was extracted from cultures using a phenol-chloroform extraction adapted from (Griffiths et al., 2000). Briefly, cells were mechanically lysed by adding to the frozen cells 0.5 g of 452-600 µm acid-washed glass beads (Sigma), 500 µL of 5 % CTAB in potassium phosphate buffer 120 mM pH 8 and 500 µL Phenol:Chloroform:Isoamyl-alcohol (25:24:1) pH 8 (Thermofisher). Tubes were then shaken for 30 s at 2500 rps with the PowerLyzer 24 (MO BIO Laboratories Inc) and centrifuged at 22,000 rcf for 10 min at 4 °C. The aqueous phase was washed with 500 µL of Chloroform:Isoamyl-alcohol (24:1) and centrifuged at 22,000 rcf for 10 min at 4 °C. 1 mL of 30% Polyethylen Glycol 6000 solution was added to the aqueous phase and incubated overnight at 4 °C to precipitate nucleic acids. Tubes were then centrifuged at 22,000 rcf for 30 min at 4 °C and pellets were washed with ice-cold 70% ethanol. Pellets were dried for 20 min at room temperature and re-suspended in 50 µL water. Genomic DNA was removed using the TURBO DNAse (Invitrogen) following the manufacturer’s instructions. RNA was precipitated for 2 hours at -20 °C with 25 µL of NaCl (6 M), 5 µL of Sodium Acetate (3 M) and 50 µL of ice-cold 2-propanol. Tubes were centrifuged, pellets were washed with ethanol and air-dried as previously described. RNA was re-suspended in a final volume of 20 µL of nuclease-free water. RNA integrity was observed using agarose gel electrophoresis and the concentration and purity were assessed by spectrophotometry (ND-1000, Nanodrop) and fluorometry (Qubit RNA High Sensitivity kit and Qubit 3, Invitrogen). None of the extraction negative controls contained detectable RNA and were thus not sent for sequencing.

Samples from the 3 RNA-seq experiments were sequenced separately. RNA extract Quality Control (QC) and sequencing were performed by Novogene (Cambridge). Quantity, purity, and integrity of the samples were assessed using Nanodrop (ThermoFisher), Agarose Gel Electrophoresis and Bioanalyzer 2100 (Agilent). Library preparation steps included rRNA depletion, RNA fragmentation, reverse transcription, Illumina adapter ligation, size selection and PCR amplification. The libraries were sequenced on the Illumina NovaSeq platform with the PE150 strategy.

### Data analysis

Data analyses were performed on R version 4.3.3 (R Core Team, 2024). All scripts used for the analysis are available as a Quarto notebook at https://doi.org/10.6084/m9.figshare.27331281.

### *In vitro* growth data

Raw OD600 values were averaged between technical replicates and the noise in OD measurement was reduced with a locally weighted scatterplot smoothing regression using the lowess function from the stats package v4.3.2 (R Core Team, 2024). Growth rates were estimated from these averaged growth curves using the growthcurver package v0.3.1 (Sprouffske & Wagner, 2016). Statistical differences between treatments for growth rates and CFU number were assessed using either a t-test, with the t.test function, or a Wilcoxon pairwise comparison when the condition of validity for the t-test were not met.

### RNA-seq data

The bioinformatic analysis was conducted on the Supercomputing Wales cluster. The reference genome and annotations from the FRB141 strain were downloaded from Genbank under the accession GCF_002291445.1 (BioProject PRJNA308225). A transcript FASTA file was produced from the reference genome and annotations using gffread v0.11.5 (Pertea & Pertea, 2020). The RNA-seq data were then analysed with the nf-core/rnaseq nextflow pipeline v3.10.1 (<u>10.5281/zenodo.1400710</u>; (Ewels et al., 2020) with raw fastq read, the reference genome, the reference annotation, and the reference transcript FASTA as input. Briefly, raw read quality was assessed with fastQC. Genome index was generated with STAR (Dobin et al., 2013) using the CDS entries from the annotation and a transcript index was generated using Salmon (Patro et al., 2017). The transcriptome reads were then aligned against the genome index using STAR and gene transcripts were quantified with Salmon and count tables were generated.

Samples from the wood tissue extract experiment (experiment 1) were then analysed on their own, and samples from the larval extract experiments (experiment 2 and 3) were analysed together. Count data were imported in R using the tximport package v1.28.01 (Soneson et al., 2016). Subsequent analyses were performed using the edgeR package v4.0.16 (Robinson et al., 2010). Transcripts with less than 10 reads in at least two samples were filtered out, read distributions were normalised using the read trimmed mean of M-values method (Robinson & Oshlack, 2010). The data was transformed for linear modelling using the voom function from the limma package v3.58.1 (Ritchie et al., 2015) and the model was fitted with the treatment variable (i.e. the identity of chemical compounds added to the culture) and the experiment batch for the larvae chemical compound experiments. Coefficients were then computed for the following contrasts: gallery wood tissue vs asymptomatic wood tissue, Ab vs water control, Ap vs water control, As vs water control, Gm vs water control, Ab vs Ap, Ab vs As, Ab vs Gm. p- values were computed and adjusted for multiple testing using the Benjamini-Hochberg method. Differentially expressed genes were determined considering a 5 % false rate discovery (adjusted p-value < 0.05) and at least a 1.5-fold change between treatment (log2 fold change (LFC) > 0.58).

## Authors contributions

SD, JV and JM had the original idea and secured funding for the study. MCC and GT designed the experiments. MC and KR performed beetle collection and rearing and set up the oak billets infestation. GT performed chemical compounds extractions and analysis, with feedback and support from JC and JV. MCC developed protocols and performed the bioactivity assays and RNA seq experiment with support and feedback from JM, JD and UH. MCC analysed the data, created the figures and wrote the first draft of the manuscript. All authors reviewed the manuscript.

## Data availability

The raw RNAseq data are available on the European Nucleotide Archive under the project accession PRJEB69286. All scripts used for the analysis are available as a computational notebook at https://doi.org/10.6084/m9.figshare.27331281, and raw data and intermediate files to reproduce the analysis are available at https://doi.org/10.6084/m9.figshare.27331488.

## Acknowledgements

The authors are grateful to Prof. Robert Jackson for his input and advice on the manuscript and to Dr Michael Birkett for checking the manuscript prior to submission.

## Supplementary material

**Table S1:** Differentially expressed genes in *B. goodwinii* cultures supplemented with gallery wood tissue extracts compared to asymptomatic wood extracts.

**Table S2:** Differentially expressed genes in *B. goodwinii* cultures supplemented with Ab larval extracts compared to water control.

**Table S3:** Differentially expressed genes in *B. goodwinii* cultures supplemented with Ap larval extracts compared to water control.

---

**Figure S1:**
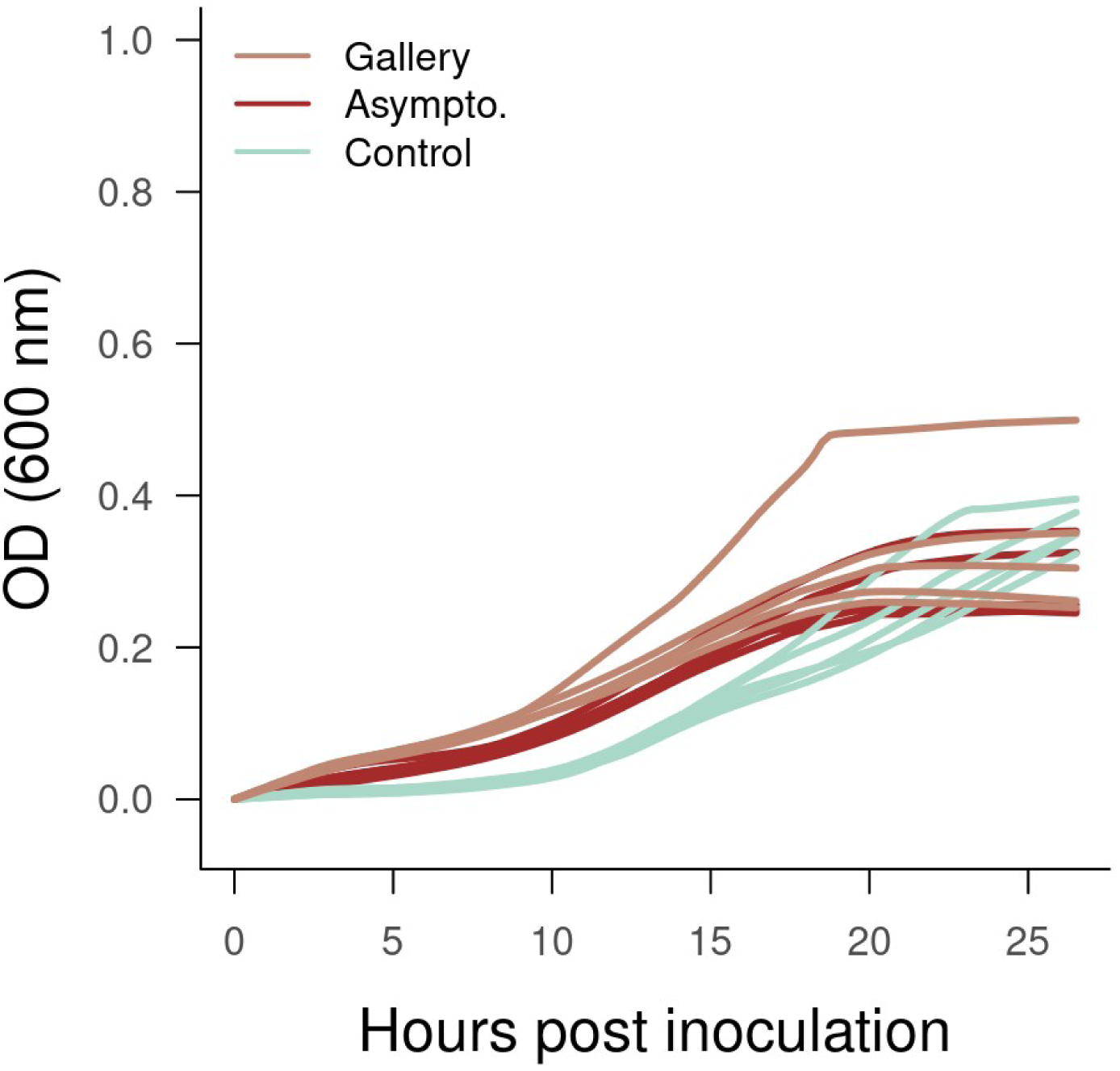
Growth curves obtained in 200 µL liquid cultures supplemented with gallery wood tissue (Gallery) and asymptomatic wood tissue (Asympto.) chemical compounds, compared to chemical compound extraction blanks (Controls).

**Figure S2:**
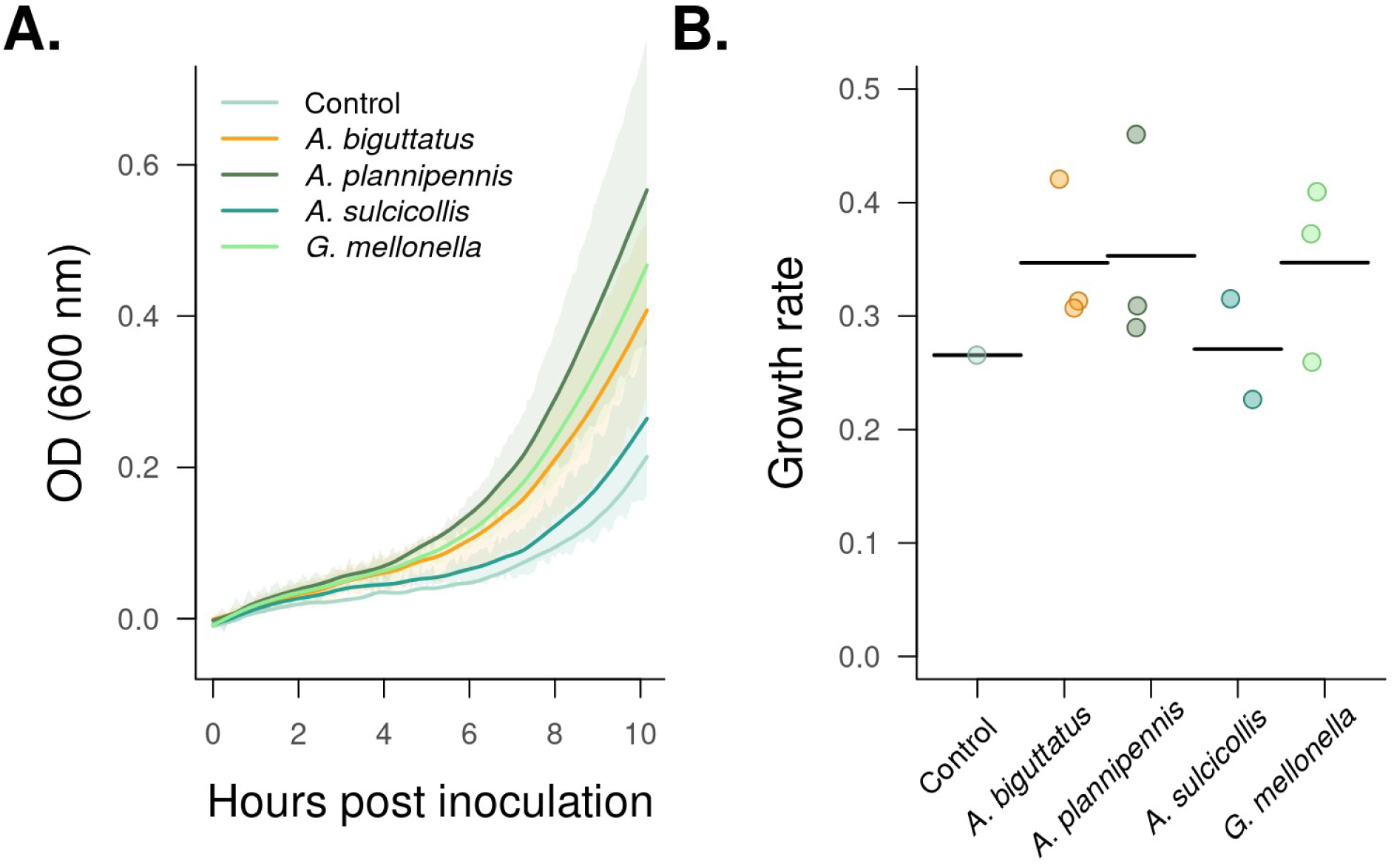
Growth of B. goodwinii cultures supplemented with insect larvae extracts. Due to the limited amount of extract, only one experiment was conducted with these larvae species and cultures were only grown for 10 hpi and collected for RNAseq analysis. **A.** OD_600_ of culture supplemented with *A. biguttatus*, *A. sulcicollis*, *A. plannipennis* and *G. melonella.* Control cultures were supplemented with extraction blanks (water). Growth was stopped at 1 hpi. **B.** Growth rates of cultures calculated on the 10 hours of growth. Differences between treatment were not significant.

## References

1. Broberg, M., Doonan, J., Mundt, F., Denman, S., & McDonald, J. E. (2018). Integrated multi-omic analysis of host-microbiota interactions in acute oak decline. Microbiome, 6(1), 21. 10.1186/s40168-018-0408-5

2. Carluccio, G., Sabella, E., Greco, D., Vergine, M., Delle Donne, A. G., Nutricati, E., Aprile, A., De Bellis, L., & Luvisi, A. (2024). Acute and Chronic Oak Decline in urban and forest ecosystems in Southern Italy. *Forestry: An International Journal of Forest Research*, cpae011. 10.1093/forestry/cpae011

3. Collmer, A., Badel, J. L., Charkowski, A. O., Deng, W.-L., Fouts, D. E., Ramos, A. R., Rehm, A. H., Anderson, D. M., Schneewind, O., van Dijk, K., & Alfano, J. R. (2000). Pseudomonas syringae Hrp type III secretion system and effector proteins. Proceedings of the National Academy of Sciences, 97(16), 8770–8777. 10.1073/pnas.97.16.8770

4. Coolen, S., Rogowska-van der Molen, M., & Welte, C. U. (2022). The secret life of insect-associated microbes and how they shape insect–plant interactions. FEMS Microbiology Ecology, 98(9), fiac083. 10.1093/femsec/fiac083

5. Correa, V. R., Majerczak, D. R., Ammar, E.-D., Merighi, M., Pratt, R. C., Hogenhout, S. A., Coplin, D. L., & Redinbaugh, M. G. (2012). The Bacterium Pantoea stewartii Uses Two Different Type III Secretion Systems To Colonize Its Plant Host and Insect Vector. Applied and Environmental Microbiology, 78(17), 6327–6336. 10.1128/AEM.00892-12

6. Davis, T. S., Crippen, T. L., Hofstetter, R. W., & Tomberlin, J. K. (2013). Microbial Volatile Emissions as Insect Semiochemicals. Journal of Chemical Ecology, 39(7), 840–859. 10.1007/s10886-013-0306-z

7. Degrave, A., Siamer, S., Boureau, T., & Barny, M.-A. (2015). The AvrE superfamily: Ancestral type III effectors involved in suppression of pathogen-associated molecular pattern-triggered immunity. Molecular Plant Pathology, 16(8), 899–905. 10.1111/mpp.12237

8. Denman, S., Doonan, J., Ransom-Jones, E., Broberg, M., Plummer, S., Kirk, S., Scarlett, K., Griffiths, A. R., Kaczmarek, M., Forster, J., Peace, A., Golyshin, P. N., Hassard, F., Brown, N., Kenny, J. G., & McDonald, J. E. (2018). Microbiome and infectivity studies reveal complex polyspecies tree disease in Acute Oak Decline. The ISME Journal, 12(2), Article 2. 10.1038/ismej.2017.170

9. Denman, S., & Webber, J. (2009). Oak declines: New definitions and new episodes in Britain. Quarterly Journal of Forestry, 103(4), 285–290.

10. Dobin, A., Davis, C. A., Schlesinger, F., Drenkow, J., Zaleski, C., Jha, S., Batut, P., Chaisson, M., & Gingeras, T. R. (2013). STAR: Ultrafast universal RNA-seq aligner. *Bioinformatics (Oxford*, England), 29(1), 15–21. 10.1093/bioinformatics/bts635

11. Doonan, J. M., Broberg, M., Denman, S., & McDonald, J. E. (2020). Host-microbiota- insect interactions drive emergent virulence in a complex tree disease. Proceedings of the Royal Society B: Biological Sciences, 287(1933), 20200956. 10.1098/rspb.2020.0956

12. Douglas, A. E. (2009). The microbial dimension in insect nutritional ecology. Functional Ecology, 23(1), 38–47. 10.1111/j.1365-2435.2008.01442.x

13. Ewels, P. A., Peltzer, A., Fillinger, S., Patel, H., Alneberg, J., Wilm, A., Garcia, M. U., Di Tommaso, P., & Nahnsen, S. (2020). The nf-core framework for community-curated bioinformatics pipelines. Nature Biotechnology, 38(3), Article 3. 10.1038/s41587-020-0439-x

14. Griffiths, R. I., Whiteley, A. S., O’Donnell, A. G., & Bailey, M. J. (2000). Rapid method for coextraction of DNA and RNA from natural environments for analysis of ribosomal DNA- and rRNA-based microbial community composition. Applied and Environmental Microbiology, 66(12), 5488–5491. 10.1128/AEM.66.12.5488-5491.2000

15. Hall, C. L., Wadsworth, N. K., Howard, D. R., Jennings, E. M., Farrell, L. D., Magnuson, T. S., & Smith, R. J. (2011). Inhibition of Microorganisms on a Carrion Breeding Resource: The Antimicrobial Peptide Activity of Burying Beetle (Coleoptera: Silphidae) Oral and Anal Secretions. Environmental Entomology, 40(3), 669–678. 10.1603/EN10137

16. Jahn, C. E., Willis, D. K., & Charkowski, A. O. (2008). The Flagellar Sigma Factor FliA Is Required for Dickeya dadantii Virulence. Molecular Plant-Microbe Interactions®, 21(11), 1431–1442. 10.1094/MPMI-21-11-1431

17. Jittawuttipoka, T., Buranajitpakorn, S., Vattanaviboon, P., & Mongkolsuk, S. (2009). The Catalase-Peroxidase KatG Is Required for Virulence of Xanthomonas campestris pv. Campestris in a Host Plant by Providing Protection against Low Levels of H2O2. *Journal of Bacteriology*, *191*(23), 7372–7377. 10.1128/JB.00788-09

18. Leroy, P. D., Sabri, A., Verheggen, F. J., Francis, F., Thonart, P., & Haubruge, E. (2011). The semiochemically mediated interactions between bacteria and insects. Chemoecology, 21(3), 113–122. 10.1007/s00049-011-0074-6

19. Malamud, F., Torres, P. S., Roeschlin, R., Rigano, L. A., Enrique, R., Bonomi, H. R., Castagnaro, A. P., Marano, M. R., & Vojnov, A. A. (2011). The Xanthomonas axonopodis pv. Citri flagellum is required for mature biofilm and canker development. Microbiology, 157(3), 819–829. 10.1099/mic.0.044255-0

20. Miller, M. B., & Bassler, B. L. (2001). Quorum Sensing in Bacteria. Annual Review of Microbiology, 55(1), 165–199. 10.1146/annurev.micro.55.1.165

21. Nomura, K., Andreazza, F., Cheng, J., Dong, K., Zhou, P., & He, S. Y. (2023). Bacterial pathogens deliver water- and solute-permeable channels to plant cells. Nature, 621(7979), 586–591. 10.1038/s41586-023-06531-5

22. Patro, R., Duggal, G., Love, M. I., Irizarry, R. A., & Kingsford, C. (2017). Salmon: Fast and bias-aware quantification of transcript expression using dual-phase inference. Nature Methods, 14(4), 417–419. 10.1038/nmeth.4197

23. Pertea, G., & Pertea, M. (2020). GFF Utilities: GffRead and GffCompare. F1000Research, *9*, ISCB Comm J-304. 10.12688/f1000research.23297.2

24. R Core Team. (2024). *R: a language and environment for statistical computing* [Manual]. R Foundation for Statistical Computing. https://www.R-project.org/

25. Reed, K., Denman, S., Leather, S. R., Forster, J., & Inward, D. J. G. (2018). The lifecycle of Agrilus biguttatus: The role of temperature in its development and distribution, and implications for Acute Oak Decline. Agricultural and Forest Entomology, 20(3), 334–346. 10.1111/afe.12266

26. Ritchie, M. E., Phipson, B., Wu, D., Hu, Y., Law, C. W., Shi, W., & Smyth, G. K. (2015). Limma powers differential expression analyses for RNA-sequencing and microarray studies. Nucleic Acids Research, 43(7), e47. 10.1093/nar/gkv007

27. Robacker, D. C., Lauzon, C. R., & He, X. (2004). Volatiles Production and Attractiveness to the Mexican Fruit Fly of Enterobacter agglomerans Isolated from Apple Maggot and Mexican Fruit Flies. Journal of Chemical Ecology, 30(7), 1329–1347. 10.1023/B:JOEC.0000037743.98703.43

28. Robinson, M. D., McCarthy, D. J., & Smyth, G. K. (2010). edgeR: A Bioconductor package for differential expression analysis of digital gene expression data. Bioinformatics, 26(1), 139–140. 10.1093/bioinformatics/btp616

29. Robinson, M. D., & Oshlack, A. (2010). A scaling normalization method for differential expression analysis of RNA-seq data. Genome Biology, 11(3), R25. 10.1186/gb-2010-11-3-r25

30. Ruffner, B., Schneider, S., Meyer, J. b, Queloz, V., & Rigling, D. (2020). First report of acute oak decline disease of native and non-native oaks in Switzerland. New Disease Reports, 41(1), 18–18. 10.5197/j.2044-0588.2020.041.018

31. Sicard, A., Zeilinger, A. R., Vanhove, M., Schartel, T. E., Beal, D. J., Daugherty, M. P., & Almeida, R. P. P. (2018). Xylella fastidiosa: Insights into an Emerging Plant Pathogen. Annual Review of Phytopathology, 56(1), 181–202. 10.1146/annurev-phyto-080417-045849

32. Singh, B. K., Delgado-Baquerizo, M., Egidi, E., Guirado, E., Leach, J. E., Liu, H., & Trivedi, P. (2023). Climate change impacts on plant pathogens, food security and paths forward. Nature Reviews Microbiology, 1–17. 10.1038/s41579-023-00900-7

33. Smee, M. R., & Hendry, T. A. (2022). Context-Dependent Benefits of Aphids for Bacteria in the Phyllosphere. The American Naturalist, 199(3), 380–392. 10.1086/718264

34. Soneson, C., Love, M. I., & Robinson, M. D. (2016). Differential analyses for RNA-seq: Transcript-level estimates improve gene-level inferences (No. 4:1521). F1000Research. 10.12688/f1000research.7563.2

35. Sprouffske, K., & Wagner, A. (2016). Growthcurver: An R package for obtaining interpretable metrics from microbial growth curves. BMC Bioinformatics, 17(1), 172. 10.1186/s12859-016-1016-7

36. Sugio, A., Kingdom, H. N., MacLean, A. M., Grieve, V. M., & Hogenhout, S. A. (2011). Phytoplasma protein effector SAP11 enhances insect vector reproduction by manipulating plant development and defense hormone biosynthesis. Proceedings of the National Academy of Sciences, 108(48), E1254–E1263. 10.1073/pnas.1105664108

37. Teh, C. H., Nazni, W. A., Lee, H. L., Fairuz, A., Tan, S. B., & Sofian-Azirun, M. (2013). In vitro antibacterial activity and physicochemical properties of a crude methanol extract of the larvae of the blow fly Lucilia cuprina. Medical and Veterinary Entomology, 27(4), 414–420. 10.1111/mve.12012

38. Thomas, D., Morgan, D. G., & DeRosier, D. J. (2001). Structures of Bacterial Flagellar Motors from Two FliF-FliG Gene Fusion Mutants. Journal of Bacteriology, 183(21), 6404–6412. 10.1128/JB.183.21.6404-6412.2001

39. Tkaczyk, M., & Sikora, K. (2024). The Role of Bacteria in Acute Oak Decline in South- West Poland. Microorganisms, 12(5), Article 5. 10.3390/microorganisms12050993

40. Troisfontaines, P., & Cornelis, G. R. (2005). Type III Secretion: More Systems Than You Think. Physiology, 20(5), 326–339. 10.1152/physiol.00011.2005

41. Volkovitsh, M. G., Orlova-Bienkowskaja, M. J., Kovalev, A. V., & Bieńkowski, A. O. (2020). An illustrated guide to distinguish emerald ash borer (Agrilus planipennis) from its congeners in Europe. Forestry: An International Journal of Forest Research, 93(2), 316–325. 10.1093/forestry/cpz024

42. Weisskopf, L., Schulz, S., & Garbeva, P. (2021). Microbial volatile organic compounds in intra-kingdom and inter-kingdom interactions. Nature Reviews Microbiology, 19(6), Article 6. 10.1038/s41579-020-00508-1

43. Wilkinson, D. A., Chacko, S. J., Vénien-Bryan, C., Wadhams, G. H., & Armitage, J. P. (2011). Regulation of Flagellum Number by FliA and FlgM and Role in Biofilm Formation by Rhodobacter sphaeroides. Journal of Bacteriology, 193(15), 4010– 4014. 10.1128/jb.00349-11

44. Worley, M. J., Ching, K. H. L., & Heffron, F. (2000). Salmonella SsrB activates a global regulon of horizontally acquired genes. Molecular Microbiology, 36(3), 749–761. 10.1046/j.1365-2958.2000.01902.x

